# Analysis of alternative splicing uncovers a vastly expanded transcriptomic response to hypoxia in human vascular endothelial cells

**DOI:** 10.64898/2025.12.09.693320

**Authors:** Justin T. Roberts, Viktor M. Pastukh, Grant T. Daly, Adeyeye I. Haastrup, Raymond J. Langley, Hank W. Bass, Mark N. Gillespie

**Author notes:** Correspondence: Justin T. Roberts, Ph.D. University of South Alabama College of Medicine Department of Pharmacology, AL 36688, Mark N. Gillespie, Ph.D. University of South Alabama College of Medicine Department of Pharmacology, AL 36688.

## Abstract

Hypoxia is a fundamental pathophysiological stimulus that plays important roles in multiple cardiopulmonary diseases. During hypoxic stress, cells adapt by undergoing widespread transcriptional reprogramming. The conventional approach to investigating this response has largely involved RNA-seq analysis which typically quantifies transcriptional output at the level of individual genes. However, as most mammalian genes encode multiple transcripts that are divergently regulated by alternative splicing, consolidating these discrete features to the gene level can obscure critical changes within the transcriptome that are largely unappreciated. To more accurately define the role of individual transcript usage during hypoxia, herein we employed three different analytical strategies that collectively identified thousands of instances of alternative splicing in human endothelial cells undergoing hypoxic stress. Notably, the set of differentially utilized transcripts displayed minimal overlap with the genes identified to be differentially expressed by conventional RNA-seq analysis, indicating divergent usage of individual transcripts does not reliably culminate in a detectable change in overall expression. This outcome is particularly clear at the earliest time-point of hypoxia where we found the transcriptional response was mediated almost exclusively by alternative splicing. A subset of these acutely responsive genes was variably detected as differentially expressed or alternatively spliced at later time points, demonstrating hypoxic transcription is highly dynamic and temporally complex. Further, the number of genes in pathways incriminated in the hypoxic response was also expanded considerably when alternatively spliced transcripts were included in the analysis, suggesting distinct attributes of the hypoxic transcriptome are unveiled by transcript level assays that would otherwise be obfuscated by standard RNA-seq analyses that summarize expression to the gene level. Collectively, these results strongly point to alternative splicing being a significant, albeit understudied, component of the transcriptional response to hypoxic stress which is vastly more complicated than previously thought.

## INTRODUCTION

High-throughput sequencing of cDNA, i.e., RNA-seq, has allowed unprecedented insight into transcriptomic pathways involved in adaptive and pathophysiologic responses to stressful stimuli. This assay primarily consists of reverse transcribing isolated RNA to generate cDNA libraries that are subsequently sequenced, making the resulting reads direct measures of individual transcripts. Paradoxically, however, most analyses of RNA-seq data collapse these transcript-resolved reads to the gene that encodes them, herein referred to as “gene level” expression. This convention arises from the assumption that gene-level counts are more accurate given the difficulty of resolving similar yet distinct transcript isoforms with short-read sequencing. Here, because the sequenced fragments are far shorter than full-length transcripts, many reads map to exons shared by multiple isoforms or to overlapping transcript structures making it technically difficult to unambiguously assign them to a single isoform. Therefore, although RNA-seq inherently provides ‘transcript level’ information, biological interpretation is still typically conceptualized at the level of genes. This approach is problematic, however, as it provides an incomplete and often misleading representation of changes in individual transcript usage, particularly when isoforms encoded by the same gene are expressed in opposing directions. In such cases, summarization to the gene level effectively masks changes in expression of individual isoforms and can support the false conclusion that the gene is transcriptionally inactive. We argue that such gene-level analytical approaches are grossly over-simplified, and that accurate profiling of transcript-level expression is required to fully illuminate cellular transcription.

The mechanism cells employ to express multiple distinct transcripts from a single gene is alternative splicing. This process, classically defined as the rearrangement of exons and introns, can also generate isoforms with variable untranslated regions (UTRs) and polyadenylation sites which mediate transcript cellular localization, stability, and regulatory capacity. The vast majority (∼95%) of human genes are alternatively spliced, effectively expanding ∼20,000 genes into more than 100,000 protein isoforms^1,2^. This marked increase in functional and structural diversity is particularly advantageous during stress conditions that require rapid cellular adaptation. Here, in contrast to proteins which are expensive to synthesize and typically persist longer, RNA transcripts are uniquely suited for quick and reversible regulation as their relatively low energetic cost and short half-life allow transcript levels to shift within minutes, thus making it an ideal substrate for temporal control. Likely the most appreciated example of this concept is the extensive reprogramming of alternative splicing that occurs in the tumor microenvironment where rapidly proliferating cells outgrow their blood supply. Here, hypoxic stress drives cancer cells to employ differential transcript usage, i.e., isoform switching, which promotes angiogenesis and vascular remodeling to support continued tumor growth. Notably, while this mechanism is well-studied in the context of cancer progression, how primary vascular cells use alternative splicing to adapt to hypoxia is not completely understood.

Accordingly, this study profiles how human vascular endothelial cells utilize alternative splicing mechanisms to facilitate transcriptional adaptation to hypoxic stress. The hypoxic transcriptome has historically been evaluated at the gene-level, and the role of alternative splicing and differential transcript usage in hypoxic adaptation has not been systemically investigated. We therefore assessed RNA-seq data in terms of splicing events and transcript usage with the aim of comparing these outputs to traditional gene level differential expression and concordant pathway analysis. Using human umbilical vein endothelial cells (HuVECs), we performed a robust transcript-level examination that assayed different attributes of alternative splicing at multiple durations of hypoxic exposure. Our analysis reveals that, in short, the hypoxic response is vastly more complicated when viewed through the lens of transcript expression.

## RESULTS

We performed RNA-seq on HuVECs cultured in normoxia and increasing durations of hypoxia (1, 3, and 24 hours). To establish a baseline for comparison with subsequent transcript-level analyses, we first assessed differential expression using conventional gene-level approaches. Only modest differences were observed between the normoxic control and the earliest hypoxic time point, with 191 genes differentially expressed. In contrast, the later time points exhibited large and distinct transcriptional shifts, with 3,495 genes differentially expressed at three hours and 4,160 at twenty-four hours (Figure 1A). Although some overlap existed among these gene sets, fewer than one percent of differentially expressed genes (n = 55) were common across conditions, and roughly half (3,549) were unique to a single hypoxic duration (Figure 1A). These results indicate that the endothelial transcriptional response to hypoxia is both extensive and highly time dependent.

**Figure 1.**
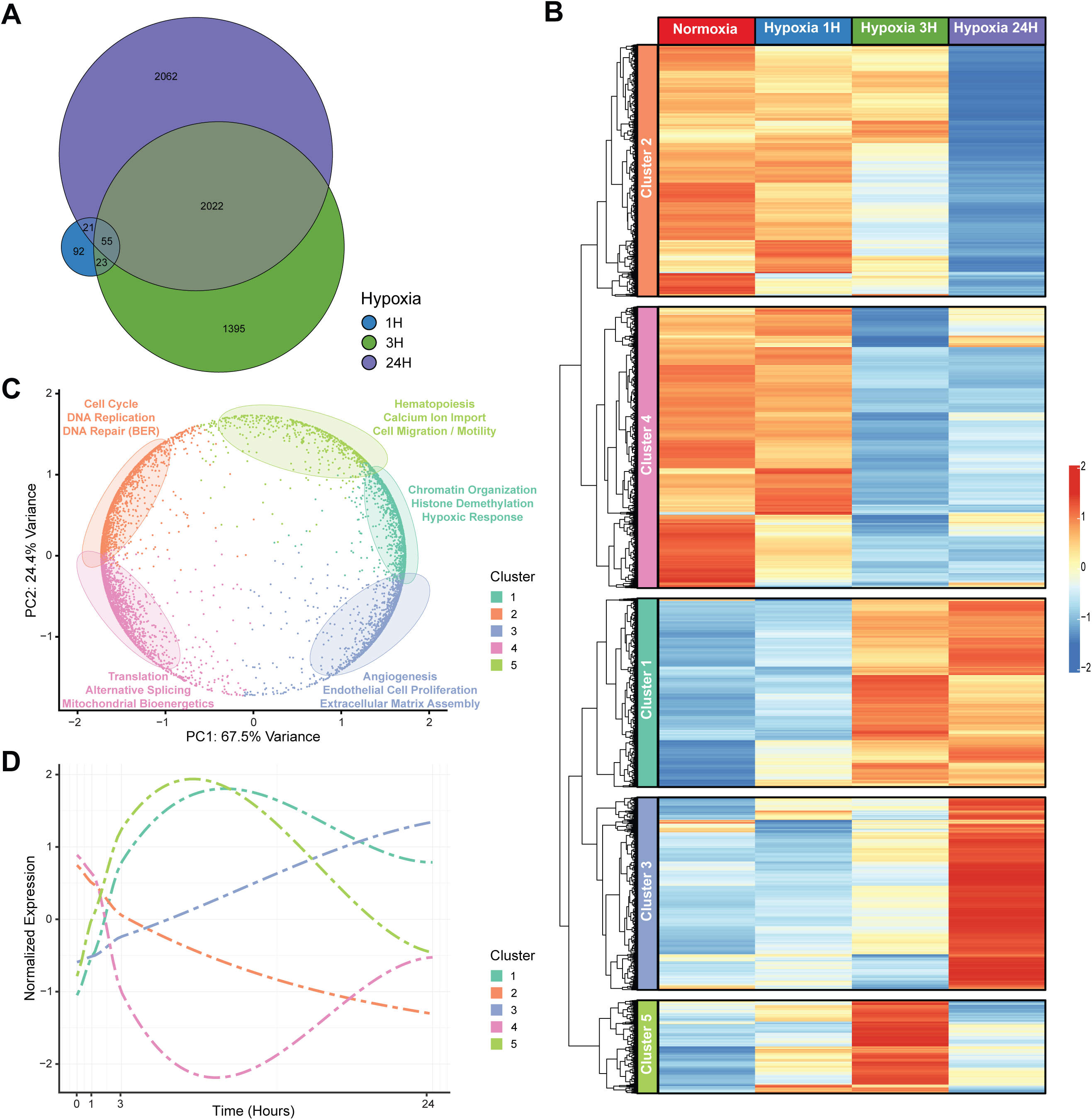
Gene-level analysis reveals coordinated and temporally structured transcriptional responses to hypoxia. (A) Venn diagram showing the overlap between differentially expressed genes (DEGs) identified at one (blue), three (green), and twenty-four (purple) hours of hypoxic exposure. (B) Heatmap showing normalized expression of significant (FDR < 0.1) DEGs identified in all hypoxic time points, clustered into five groups based on shared temporal expression patterns. Columns represent the four experimental conditions, with replicate samples averaged and indicated by condition-specific color blocks. Rows represent individual DEGs and are split into clusters and indicated by a colored side bar. Heatmap values represent row-scaled z-scores (−2 to 2), with warmer colors indicating higher relative expression and cooler colors indicating lower expression. (C) Principal component analysis (PCA) plot of the five identified gene clusters. Each point represents a gene, positioned according to its expression profile across the hypoxic time course, with cluster membership indicated by color. Enriched biological processes (as determined by gene ontology analysis) are shown alongside the corresponding set of genes, highlighting the major functional pathways represented in each group. Ellipses represent the 80% confidence region for each cluster, and the variance of each principal component is indicated on the y-axes (as determined by PCA). (D) Plot depicting LOESS-smoothed temporal expression trajectories for each of the identified clusters. Lines (colored according to cluster assignment) represent fitted curves interpolating between the four measured time points, showing the mean normalized expression (y-axis) of all member genes at each hypoxic duration (x-axis).

To further characterize the observed temporal structure, we performed clustering analysis based on expression dynamics over time. This revealed five discrete groups of genes exhibiting transcriptional patterns specific to individual hypoxic time points (Figure 1B). Gene ontology (GO) enrichment analysis showed that each cluster corresponded to a coherent set of biological processes associated with hypoxic adaptation (Figure 1C). Cluster 1 comprised genes consistently upregulated at the later time points and enriched for pathways such as chromatin organization and histone demethylation that are hallmarks of canonical hypoxic signaling^3,4^. Cluster 2 encompassed genes strongly downregulated at twenty-four hours and involved in cell cycle regulation, DNA replication, and DNA repair, aligning with prior observations that prolonged hypoxia induces cell cycle arrest and suppression of DNA synthesis through activation of cyclin-dependent kinase inhibitors^5–9^. Cluster 3, also specific to twenty-four-hour hypoxia, featured genes showing strong induction in pro-angiogenic pathways such as cell proliferation and extracellular matrix assembly that are critical for new vascularization^10–12^. Cluster 4 included genes exhibiting decreased expression at both later time points and involved in alternative splicing, translation, and mitochondrial bioenergetics, all processes known to be reprogrammed during sustained hypoxia^13–19^. Cluster 5 contained genes that transiently increased at three hours and associated with hematopoiesis, calcium import, and cell migration/motility, all key contributors to neovascularization and angiogenesis^20–25^. Together, these results demonstrate that the endothelial hypoxic transcriptome is governed by coordinated, temporally organized shifts in gene expression (Figure 1D). The major functional modules identified also align with well-established components of the hypoxic response, therefore validating our experimental and analytical approach and providing a robust framework for subsequent comparison with the transcript-level assays.

Having established the transcriptional response to hypoxia at the gene level, we next extended our analysis to the level of individual transcripts by examining the same RNA-seq time course for isoform-specific changes. To investigate this as rigorously as possible, we employed three complementary algorithms that quantify condition-dependent differences in distinct attributes of alternative splicing: DEXSeq^26^, which assesses differential exon usage by counting reads that align to individual exons; DRIMSeq^27^, which evaluates shifts in relative transcript abundance within genes; and SUPPA2^28^, which detects specific splicing events (e.g., exon skipping, intron retention) from junction-spanning reads (Figure 2A). Application of these approaches revealed substantial differences in the number and identity of alternatively spliced genes detected at each time point (Figure 2B). Across all durations of hypoxic exposure, DEXSeq identified the largest number of alternatively spliced genes — approximately 2–3 times more than DRIMSeq and SUPPA2 combined — consistent with its higher sensitivity arising from testing many more features per gene, whereas SUPPA2 detected an intermediate number of events and DRIMSeq reported the fewest. Notably, overlap among the methods was limited, with fewer than five percent of alternatively spliced genes at any time point identified by all three tools, indicating that each algorithm captures a largely non-redundant component of transcript-level regulation.

**Figure 2.**
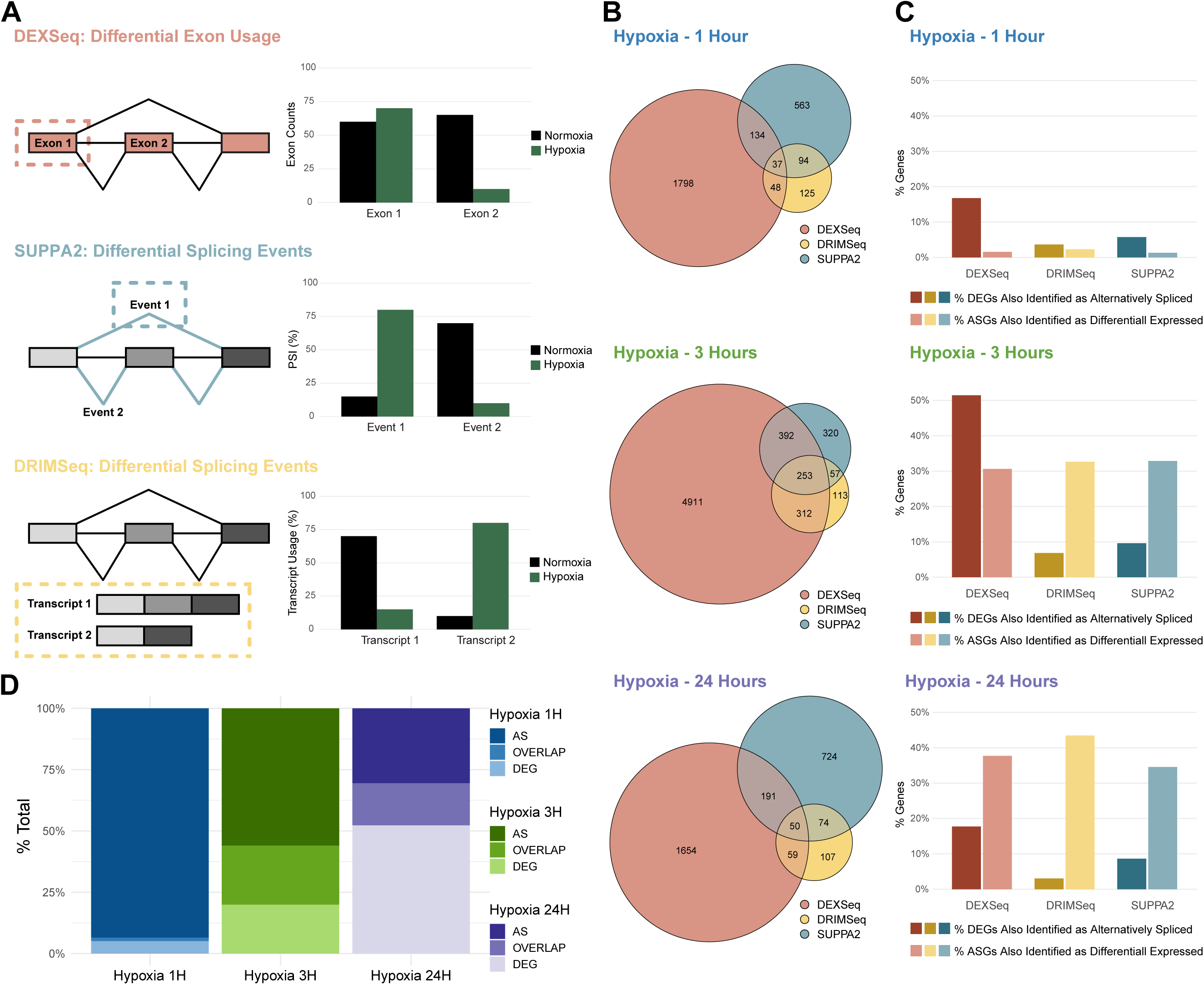
Transcript-level analysis uncovers extensive and temporally dynamic alternative splicing profiles that are largely uncaptured by gene-level approaches. (A) Schematic overview of the three algorithms used to quantify changes in alternative splicing. Exons are shown as grayscale rectangles and splicing events as connecting lines. The features evaluated by each method are highlighted with tool-specific colors and dotted outlines (DEXSeq – individual exons, light red; DRIMSeq – full transcript models, gold; SUPPA2 – specific splicing events, teal). For each method, an accompanying sample plot illustrates how condition-dependent changes of the relevant feature (x-axis) would be quantified, with y-axes showing the respective values as indicated (PSI, percent spliced-in). (B) Venn diagrams showing the overlap of alternatively spliced genes detected by DEXSeq (light red), DRIMSeq (gold), and SUPPA2 (teal) at each indicated hypoxic time point. (C) Bar plots comparing alternatively spliced genes reported from each algorithm to the set of differentially expressed genes (DEGs) previously identified. For each method, dark bars represent the percentage of total DEGs also identified as alternatively spliced, and light bars represent the percentage of alternatively spliced genes that were also classified as DEGs. Colors correspond to DEXSeq (dark/light red), DRIMSeq (dark/light gold), and SUPPA2 (dark/light teal). The y-axes indicate the percentage of genes in each category. (D) Stacked bar plots summarizing, for each hypoxic time point (x-axis), the relative proportions of genes identified as alternatively spliced (AS), differentially expressed (DEG), or detected by both (OVERLAP). The y-axis shows the percentage of total genes represented by each category. Colors as indicated in the plot.

Comparing these results to the differentially expressed genes (DEGs) identified at each time point revealed striking temporal changes in how transcript-level regulation aligns with gene-level expression (Figure 2C). At all durations of hypoxia, DEXSeq exhibited the highest overlap with DEGs, whereas the two non-exon-centric methods consistently showed much lower concordance. At one hour, overlap was minimal: DEXSeq identified ∼17% of DEGs as alternatively spliced, and DRIMSeq and SUPPA2 each captured fewer than 6%, with only ∼1–2% of alternatively spliced genes also detected as differentially expressed. In contrast, at the three and twenty-four-hour time points, transcript-and gene-level responses showed much stronger convergence. Although the fraction of DEGs also detected as alternatively spliced by DRIMSeq and SUPPA2 remained low (<10%), approximately one-third (∼30–40%) of alternatively spliced genes identified by any method were also differentially expressed. DEXSeq showed a stronger but time-dependent correspondence in the fraction of DEGs detected as alternatively spliced, identifying more than half at three hours but returning to earlier levels at twenty-four hours. These patterns likely reflect the dramatic expansion in the number of DEGs identified during the longer hypoxic exposures, which increased the absolute overlap between the two regulatory layers.

Grouping all alternatively spliced genes reported by the three methods together and comparing them to the total set of identified DEGs at each time point recapitulated these temporal trends and underscored the limited overall concordance between transcript-and gene-level responses (Figure 2D). Notably, across the time course, only ∼15% of genes were shared between the two analyses. Acute hypoxia was dominated by splicing changes that showed minimal accompanying differences in gene expression, with only ∼1% of alternatively spliced genes also detected as DEGs, indicating that the earliest response to hypoxic stress is largely at the transcript level and would be overlooked by gene-level analysis alone. At the intermediate duration, overlap between DEGs and alternatively spliced genes was most pronounced, reflecting the strongest integration of expression changes and splicing regulation. By twenty-four hours, the transcriptional response shifted more towards a profile dominated by gene-level changes, yet still retained substantial transcript-level regulation, indicating that prolonged hypoxia engages both these processes in parallel. Together, these findings demonstrate that analysis of alternative splicing uncovers a significant and temporally dynamic component of the hypoxic response that remains largely invisible to conventional gene-level approaches.

Disparities in temporal expression at the gene-level coupled with minimal consensus between alternatively spliced and differentially expressed genes prompted us to next assess how all our data generated thus far collectively converged. By comparing results from each analysis, we determined that despite the large size of datasets, genes identified by the two analytic strategies were relatively distinct at each time point (Figure 3A). Of the top six largest gene sets identified, four were unique to one analysis or time point, and only seven genes in total were common to all six datasets. The highest degrees of overlap were evident at the later durations of hypoxic exposure for both gene-and transcript-level detection, indicating a subset of genes maintained differential expression or alternative splicing throughout these time points. Intrigued by the large disparities in the total number of genes identified as alternatively spliced versus differentially expressed, we wondered whether the same genes toggled between the two regulatory layers over the course of hypoxic exposure. Because the predominant transcriptional activity of hypoxia after one hour was primarily identified as alternative splicing, we tracked the fate of transcripts detected at this time point. We found that genes spliced at one hour were variably detected as differentially expressed or as alternatively spliced at the later time points; for instance, ∼50% of genes alternatively spliced at one hour are also spliced at three hour and then have no expression at 24 hour (Figure 3B). This observation implies that isoform selection, which occurs throughout the hypoxic transcriptional response, may undermine the ability of RNA-seq analysis to comprehensively characterize the hypoxic transcriptome.

**Figure 3.**
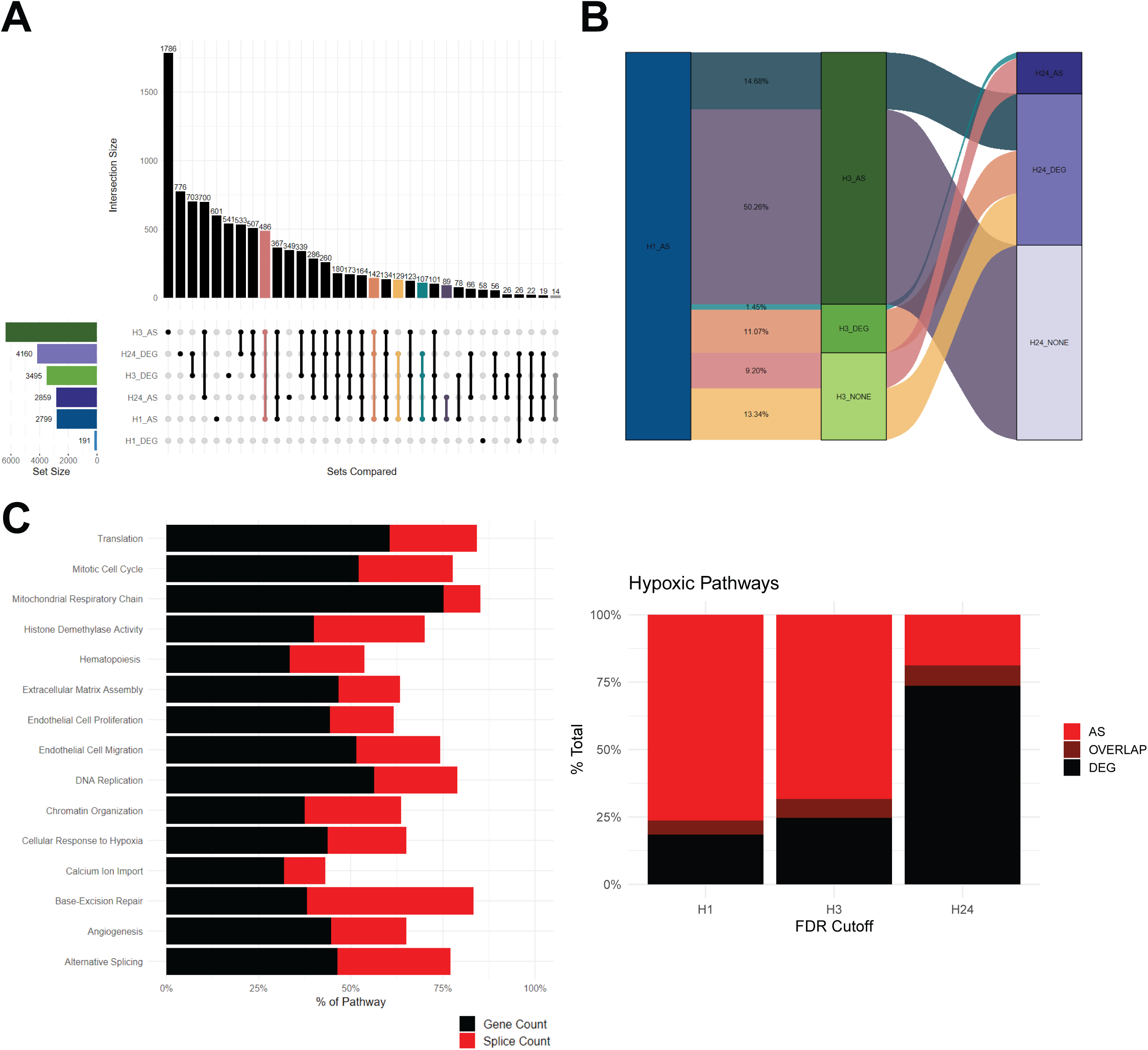
Analysis of alternative splicing provides additional insight into the functional response to hypoxia. (A) Upset plot illustrating overlap among differentially expressed (DEGs) and alternatively spliced (AS) genes identified at each hypoxic time point. Bars (top) represent the total number of genes (also labeled above each bar) belonging to each intersection (bottom), denoted by connected dots between each set of genes. Bars (left) represent the number of genes in each indicated set, colored by condition. (B) Alluvial plot illustrating the temporal fate of genes identified as alternatively spliced at one hour of hypoxia (H1_AS). Each ribbon represents a group of genes and tracks whether they are subsequently detected as alternatively spliced (AS), differentially expressed (DEG), or unchanged (NONE) at three (H3) and 24 (H24) hours. Ribbon widths are proportional to the number of genes in each category, with the specific percentage of the total indicated. (C) Stacked bar plots showing the contribution of gene-level (black) and transcript-level (red) results to the percentage of genes contained in the 15 biological pathways previously identified to be enriched in hypoxia (left), and the percentage of total enriched pathways (y-axis) detected by each analysis, labeled for each hypoxic time point (x-axis) and colored based on what level of analysis the pathway was detected by (right).

To determine if integrating transcript-level information with conventional gene-level analyses affords new insights adaptive response to hypoxia, we asked whether the pathways identified using gene-level analysis also included genes revealed by alternative splicing analysis. This analysis revealed that total representation within each pathway increased substantially transcript-level data was incorporated (Figure 3C). Subsequent GO analysis performed using the total set of alternatively spliced genes at each time point further showed that the aggregate number of detected pathways expanded markedly relative to the previous gene-level results, specifically at one-and three-hour hypoxia. Notably, overlap between the two ontology analyses was minimal, suggesting that the transcript-level data is capturing unique and previously undetected functional attributes of the hypoxic response. Collectively, these results demonstrate that including transcript-level data into pathway analyses results in a considerable gain beyond that obtained by gene-level analysis alone.

To understand how individual genes can be variably detected as either alternatively spliced or differentially expressed, it is useful to consider how differential isoform usage impacts detection of transcriptional activity at the level of genes. For example, if individual transcripts encoded by the same gene are expressed in opposite directions, they may effectively offset each other such that the changes in each transcript, when summarized to the gene level, fail to reach the threshold for detection as a differentially expressed gene. Alternatively, if the aggregate change in isoform switching, regardless of the direction of individual transcript expression, exceeds the threshold for detection at the gene level, then summarizing these aggregate changes will be identified as a differentially expressed gene. To explore this concept, we used an algorithm, IsoformSwitchAnalyzeR^29^, this is specifically designed to detect instances of isoform switching. Hundreds of isoform switches were identified at each hypoxic time point (Figure 4A). Depicting differential isoform usage as a function of expression assessed at the gene level showed conspicuous time dependent differences. For example, whereas there were few isoform switches detected as DEGs at one hour, at later points there was roughly equal number also identified at the gene level. Notably, while roughly half of the switches identified at each hypoxic time point were also identified as differentially expressed, the remaining half did not result in observed changes in gene expression (Figure 4B).

**Figure 4.**
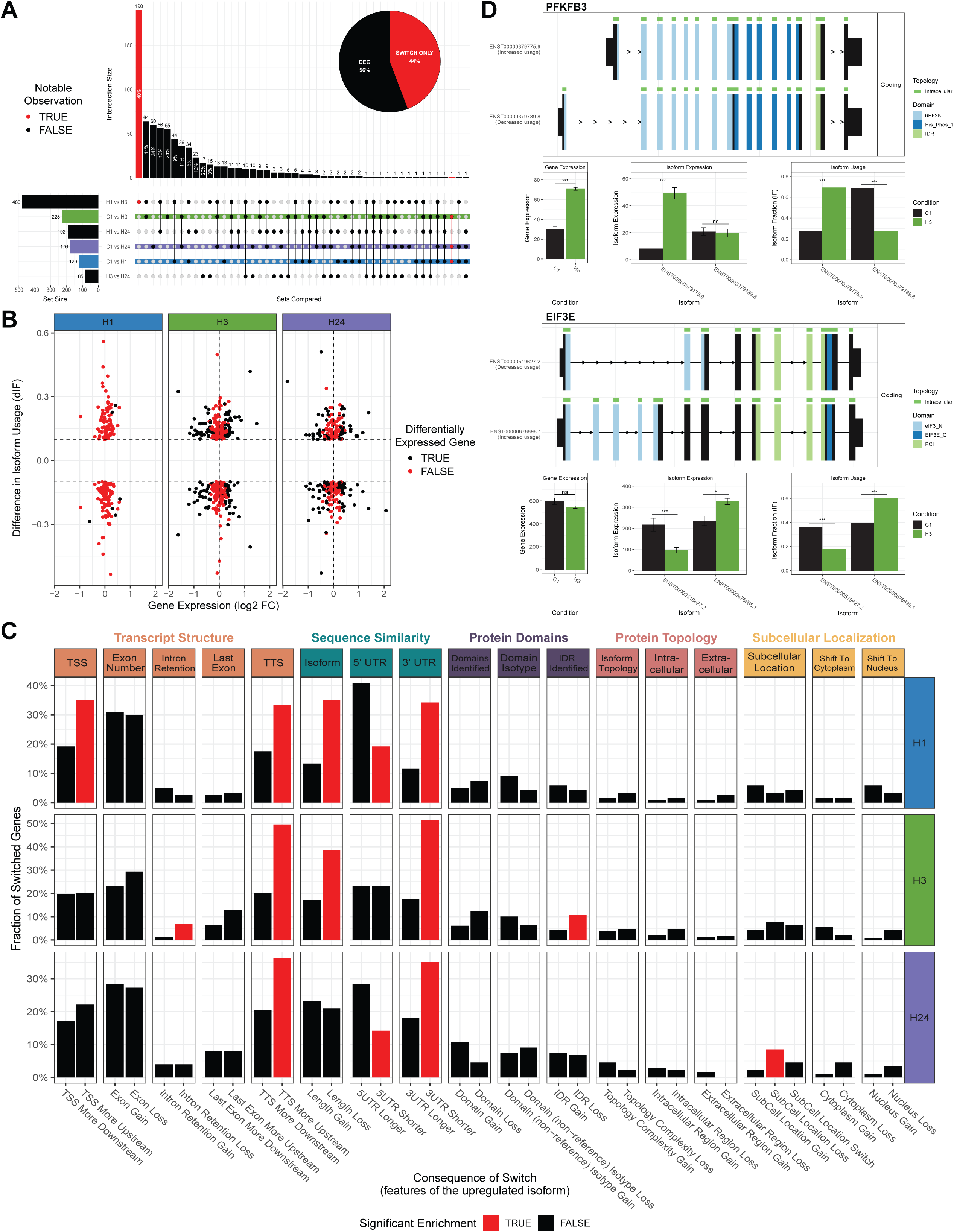
Analysis of alternative splicing provides additional insight into the functional response to hypoxia. (A) Upset plot summarizing the overlap of identified isoform switching events across all pairwise conditions. Horizontal bars (black) indicate the size of each individual switch set, and vertical bars show the size of each intersection, labeled with both the number of genes and their percentage of all switching genes. The dot matrix below depicts which conditions are included in each intersection, and selected intersections and sets are highlighted with colored stripes and points to emphasize notable observations discussed in the text. The inset pie chart summarizes the proportion of switching genes relative to normoxia that are also differentially expressed at the gene level (red) versus those regulated primarily at the isoform level without significant gene-level changes (black). (B) Dot plot illustrating the relationship between gene-level expression changes and isoform switching events identified at each hypoxic time point. Each point represents an isoform switch, with the corresponding gene-level log₂ fold change (x-axis) plotted against the change in isoform usage (dIF, y-axis). Panels are faceted by hypoxic time point, and points are colored according to whether the associated gene is also significantly differentially expressed (black) or not (red). Vertical dashed lines denote no change in gene expression (log₂FC = 0), and horizontal dashed lines indicate |dIF| = 0.1, marking switches with substantial shifts in isoform usage. (C) Bar plots summarizing predicted functional consequences of the identified isoform switches in each hypoxic time point. For each duration (rows; colored by condition), bars show the fraction of switched genes exhibiting a given consequence (columns; grouped and colored by functional category), including changes in transcription start/termination sites (TSS/TTS), 5′/3′ untranslated regions (UTR), and intrinsically disordered regions (IDRs). The y-axis indicates the fraction of switched genes within each category, and bars are colored based on whether the predicted consequence is significantly enriched relative to the opposite outcome (red) or not (black).

Conventional understanding holds that the primary consequence of isoform switching is the production of functionally distinct protein variants, however its role in hypoxic transcription seems to be more expansive. Our analysis reveals that the major impact of using different isoforms does not concern protein function *per se*, but rather, the intrinsic properties of the mRNA transcript itself (Figure 4C). For example, two common outcomes of DTU across all three hypoxic time points are alteration in the lengths of 5’ and 3’ UTRs and increased usage of upstream of transcript termination sites (TTSs). We also found that some distinguishing features are unique to specific time points. At one hour, for example, there is increased usage of alternative transcription start sites (TSSs), while at twenty-four hours there is loss of subcellular localization signals governing targeting to the nucleus or cytoplasm. Collectively, differential transcript usage in hypoxia seems critical to control not only the protein isoforms produced, but other aspects of transcript structure that confer specific regulatory properties.

## DISCUSSION

While RNA-seq remains the standard strategy for assessing transcriptional changes between experimental conditions, the resulting data are typically summarized to the gene level and therefore often interpreted as a measure of *gene* expression. However, this simplification overlooks the fact that *transcripts*, not genes, are the true functional units of expression in cells. Stated differently, whereas genes define the regulatory DNA loci that are bound by transcription factors, the actual mediators of transcription are the resulting RNA transcripts, many of which are expressed as multiple distinct isoforms. Against this background, the major finding of this study is that differential usage of such transcripts constitutes a major component of transcriptional regulation, yet this information is largely lost during conventional gene-level RNA-seq analysis.

To investigate this finding in a physiologically meaningful context, we focused on hypoxia, a fundamental microenvironmental stimulus that profoundly influences lung biology and disease. Although alternative splicing has previously been implicated in hypoxic adaptation^19,30,31^, to our knowledge this work represents the first systematic comparison of hypoxia-induced transcriptional changes assessed at both the gene and transcript levels. Notably, we found that while gene-level analyses recapitulated canonical hypoxia-responsive pathways and expression patterns, transcript-level assays substantially expanded the number of regulated loci and uncovered unique molecular processes and temporal dynamics that refine our understanding of the hypoxic regulation. Our results therefore demonstrate that conventional gene-level summarization obscures critical aspects of hypoxic transcription and that accurate conceptualization of cellular adaptation to low oxygen requires analyses that preserve isoform-level resolution.

To robustly define the complete set of genes undergoing alternative splicing during hypoxia, we employed multiple orthogonal algorithms that interrogate complementary aspects of RNA processing. Collectively, these methods span analytical perspectives and differ not only in statistical design but also in the distinct mechanistic features they are optimized to detect. DEXSeq and SUPPA2, for instance, are highly sensitive to localized splicing changes such as exon and splice-junction selection, but because they quantify individual events rather than complete isoforms, their outputs cannot always be directly linked back to the full transcript structures from which those events arise which can complicate validation. In contrast, DRIMSeq evaluates differential transcript usage within each gene, providing a more integrative view of isoform architecture and function that is less likely to track with gene-level expression, albeit with reduced sensitivity to more subtle splicing variations relative to the other tools. Using all three approaches therefore allowed us to balance mechanistic resolution with biological interpretability and achieve a more broad and comprehensive determination of the adaptive transcriptional response to hypoxia. In practice, these strategies yielded variable gene sets with modest overlap, indicating that each captures a distinct facet of transcriptional complexity rather than a redundant signal. However, all the methods consistently revealed numerous alternatively spliced genes that were absent from gene-level analyses, raising the question of why the gene-and transcript-level responses to hypoxia diverge so markedly.

Intuitively, one might expect substantial overlap between genes identified by differential expression and those flagged as alternatively spliced, with transcript-level analyses simply refining gene-level results. However, the limited concordance observed between these two levels of detection likely reflects the fact that they probe fundamentally distinct biological dimensions of transcriptional regulation that are not necessarily coupled – just as a gene may undergo strong induction or repression without altering isoform composition, it may also extensively rewire isoform balance without changing total expression. From a biological standpoint, this minimal overlap is informative and reveals overall transcriptional output and splicing regulation function as partly independent layers of the hypoxic response. At our earliest hypoxic timepoint, for instance, few changes were detected at the gene level relative to thousands at the transcript level, indicating that alternative splicing is the predominant transcriptional adaptation during acute hypoxia. Here, isoform-level regulation provides a rapid and flexible means of transcriptome remodeling, allowing cells to fine-tune regulatory protein domains or engage surveillance pathways such as nonsense-mediated decay without requiring gene-level changes in abundance. At later time points, global and isoform-specific responses became more balanced but still exhibited limited overlap, suggesting these processes can also act in parallel to shape the response to prolonged hypoxic exposure.

While these findings underscore the complexity of how hypoxia remodels the transcriptome across multiple regulatory tiers, several technical and analytical factors may also have influenced the degree of overlap observed between gene-and transcript-level results. Because methods that quantify alternative splicing rely on well-defined transcript annotations, the incompleteness of current reference data can disproportionately suppress detection of isoform-level changes relative to gene-level abundance estimates which are comparatively tolerant to such omissions. While we attempted to minimize this issue by using a common annotation across all analyses, the inherent limitations of existing transcript models likely still reduced the observed concordance. We also intentionally omitted fold-change cutoffs in the gene-level analysis, which increased the number of hypoxia-responsive genes but provided a more permissive framework for comparison to the transcript-level. Conversely, splicing algorithms typically apply more stringent significance thresholds and filter out lowly expressed isoforms which limit the set of testable genes and thus constrain opportunities for overlap. To control for such analytical variation, we standardized filtering and threshold criteria across tools to ensure comparable technical conditions and examined how varying parameters such as read quantification and reference-level selection influenced detection sensitivity. Despite these efforts, the broader trends remained unchanged as both gene-and transcript-level approaches continued to identify largely distinct subsets of hypoxia-responsive genes. Thus, the limited concordance we observed is unlikely to arise from technical artifacts but instead reflects genuine biological divergence. Collectively, the persistent lack of consensus under permissive and harmonized conditions underscores that transcript-level analyses capture distinct and biologically meaningful dimensions of the hypoxic response that are effectively masked by gene-level summarization.

To exemplify how isoform-resolved analysis can either reinforce or substantially extend interpretations drawn from gene-level data, we highlighted two genes relevant to hypoxia that illustrate each regulatory pattern. PFKFB3, representative of a gene captured at both the gene and transcript levels, encodes one of four tissue-specific isoenzymes that regulate glycolytic flux by converting fructose to stimulate glycolysis. Given that hypoxia broadly enhances glycolysis to sustain ATP production under reduced oxygen, PFKFB3 is a well-established hypoxia-inducible gene and direct transcriptional target of HIF-1. Our findings confirm the canonical upregulation at the gene-level across all hypoxic time points and further show that hypoxia induces a clear isoform switch, wherein the transcript expressed during hypoxia differs from that predominant in normoxia. Although the overall exon composition of these isoforms is largely similar, the primary difference is the use of an alternative transcription start site (TSS) in the hypoxic isoform. This is notable given that alternative TSS usage is a major mechanism of transcript diversification found in the majority of human genes ^32–34,34–37^, allowing the expression of isoforms with variable 5′ sequences that can affect transcript stability^38,39^, translational activity^40,41^, and subsequent selection of internal splice sites^42,43^ and 3’ alternative polyadenylation^44^. In short, the selection of TSSs is a key regulator of RNA processing that can directly dictate the expression of specific isoforms^45^, thereby assisting in the cellular response to environmental stress^46^. In the case of PFKFB3, this shift may facilitate rapid transcriptional activation or alter post-transcriptional regulation under hypoxic stress, adding another layer of complexity to how this key metabolic enzyme contributes to cellular adaptation. More broadly, such examples illustrate how isoform-level regulation can refine well-characterized hypoxic pathways, revealing subtle but functionally relevant dimensions of gene control that are missed by gene-level analyses alone.

In contrast to PFKFB3, the EIF3E gene illustrates how isoform-level regulation can occur independently of overall gene expression. EIF3E encodes a core subunit of the eukaryotic translation initiation factor 3 (eIF3) complex which coordinates mRNA recruitment to the ribosome and thereby modulates both global and transcript-selective translation. Our analysis revealed a hypoxia-induced isoform switch where a transcript containing additional 5′ exons and an extended 5′ UTR was enriched relative to the canonical normoxic isoform, thereby extending the N-terminal region that harbors binding domains known to mediate protein–protein interactions within the initiation complex. Inclusion of these hypoxic exons therefore likely alters the structural interface through which EIF3E engages other eIF3 subunits, potentially modifying translational selectivity or efficiency under stress. The usage of a slightly different 5′ UTR in the hypoxic isoform further implies an additional layer of regulation at the level of translation initiation, as variation in 5′ structure can modulate ribosome scanning, mRNA stability, and responsiveness to cellular stress. Notably, EIF3E has also been shown to interact directly with HIF-2α and to influence hypoxia-responsive translation by recruiting HIF-regulated transcripts to ribosomes, suggesting that isoform-specific changes could fine-tune this interaction and selectively influence the translation of hypoxia-induced mRNAs. Collectively, these observations position EIF3E as a representative case where hypoxia qualitatively remodels transcript architecture rather than evoking quantitative changes in expression, highlighting the unique biological insight revealed through isoform-resolved analysis.

Although short-read RNA-seq, as demonstrated herein, continues to serve as an effective tool for profiling global expression patterns, its capacity to resolve isoform-level changes is constrained by the inherent disconnect between read length and transcript structure. In contrast, emerging long-read sequencing technologies (e.g., Oxford Nanopore and PacBio) provide a notable improvement by directly sequencing full-length transcripts, thereby circumventing these limitations. These approaches not only enhance isoform detection and quantification but will also likely refine transcript annotations as their use becomes more widespread, which should, in turn, mitigate much of the discordance between gene-and transcript-level analyses observed in this study. Given that much of the hypoxic response appears to occur through isoform-specific regulation, this added resolution will be especially valuable for future studies. As these technologies continue to improve in accuracy and throughput, their routine incorporation will become essential for fully resolving how stressors like hypoxia rewire the transcriptome at the level of individual isoforms, a dimension of regulation that, as our results demonstrate, remains partly obscured in conventional short-read analyses summarized to the gene level.

## METHODS

### Cell culture and RNA isolation

HUVECs (ATCC, Cat. No. PSC-100-010) were cultured in endothelial growth media under standard culture conditions (5% CO2, 95% room air, 37°C) and maintained between passages 5–7. Approximately 900,000 cells were seeded into 100 mm culture dishes and typically reached 90–95% confluency within 24–48 hours. Cells were certified as mycoplasma-free by the supplier; STR genotyping was not performed. At 90–95% confluency, cells were subjected to hypoxia (2% O2) for 1, 3, or 24 hours. Following hypoxia treatment, total RNA was isolated from cells using PrepEase RNA Spin Kit (USB) according to the manufacturer’s protocol. RNA quality and concentration were assessed using a NanoDrop spectrophotometer. Samples were then shipped on dry ice via overnight delivery to Novogene (Sacramento, CA, USA) for sequencing.

RNA sequencing of triplicate samples harvested from each of the four conditions was performed by Novogene on a fee-for-service basis. Briefly, poly-T oligo magnetic beads were used to isolate messenger RNA (mRNA) from total RNA, which was then fragmented to lengths of 150–350 bases. The mRNA was reverse-transcribed to generate a cDNA library which was quantified using a Qubit fluorometer, and the size distribution was verified with a Bioanalyzer. Libraries were pooled and sequenced on an Illumina platform to generate paired-end reads.

### Data processing and differential expression analysis

To generate transcript-level abundance estimates for differential gene expression and alternative splicing analyses, reference genome and transcript annotations were obtained from the GENCODE v45 database and the RNA-seq data for each condition was quantified using Salmon^47^ (v1.10.2). The resulting transcript-level estimates were imported into R (v4.3.1) and summarized to the gene level using tximport^48^. Differential expression analysis was then performed using the standard DESeq2 (v1.42.0) workflow, with samples labeled by experimental condition. Based on principal component analysis, one replicate from the 24-hour hypoxia condition was excluded from downstream analysis. Genes with a significant adjusted p-value (<0.10) were considered differentially expressed. For visualization, normalized counts were averaged across biological replicates for each condition and row-scaled prior to heatmap generation.

### Clustering and functional analysis

To identify temporal expression patterns among hypoxia-responsive genes, we performed unsupervised clustering on the normalized gene counts. We evaluated the optimal number of clusters using several complementary approaches, including internal stability metrics (*clValid*), NbClust-based heuristics, and gap-statistic estimation. Based on these evaluations, k-means clustering (k = 5) was applied to partition genes according to shared expression trajectories across hypoxic time points. Cluster assignments were visualized using PCA and later used for heatmap annotation. To characterize the biological functions associated with each identified cluster, Gene Ontology (GO) over-representation analyses was conducted using the PANTHER database, with all hypoxia-responsive genes serving as the background reference. The resulting enrichments were compiled and exported for downstream visualization.

### Alternative splicing analysis

To characterize hypoxia-induced changes at the transcript level, we employed three complementary approaches that assay different attributes of alternative splicing. First, differential transcript usage (DTU) was assessed using DRIMSeq. Briefly, transcript-level count tables were used to fit to a Dirichlet-multinomial model of expression and tested for DTU between the conditions. To control the family-wise error rate, we applied stageR^49^ in a two-stage hierarchical testing framework to generate FDRs used for filtering (<0.10). The resulting gene sets and their regulated transcripts from each hypoxic condition were exported for downstream integration with the gene-level results. Second, exon-level differential usage was assessed with DEXSeq on STAR-aligned BAM files. GENCODE v45 annotation was converted into flattened Breifly, exon-level features were extracted from the transcript annotations using scripts supplied by the algorithm, restricting exonic parts to single genes. Read counts overlapping exonic parts were obtained from paired-end BAMs for each hypoxic condition, and differential exon usage was assessed by exon-specific dispersion and GLM expression models fitted with multi-core processing. Significantly regulated exons (FDR < 0.10) were calculated using DEXSeq-specific methods. Third, event-level alternative splicing events were quantified using SUPPA2. Using a precomputed event annotation file generated from the same annotations, percent-spliced-in (PSI, Ψ) values were calculated for each alternative splicing event type across all conditions. Differential splicing analyses were performed with Python scripts supplied by the algorithm, providing a filtered set of significantly regulated events used for integration with the results from the other tools.

### Isoform switch analysis and predicted functional consequences

Isoform switching between conditions was quantified and functionally annotated using IsoformSwitchAnalyzeR. Transcript-level counts from each condition were used to build a list of isoform switching events, restricting analyses to isoforms with full-length open reading frames. Differential isoform usage between conditions was assessed and summarized per gene. To characterize the potential functional impact of isoform switching, we extracted coding sequences and protein sequences from the identified events and subjected them to a panel of external analysis tools (CPC2 – coding potential; Pfam – protein domains; SignalP 6 – signal peptides; IUPred2A – intrinsically disordered regions; DeepTMHMM – transmembrane topology; DeepLoc2 – subcellular localization. We also classified the underlying splicing event types and compared open-reading frame usage. These analyses were used to predict and summarize the functional consequences of isoform switches on features such as coding sequence and UTR similarity, exon number, intron retention, protein domain content, intrinsically disordered regions, membrane topology, and predicted subcellular localization. For downstream interpretation, we summarized the fraction of switched genes associated with each consequence category and tested for enrichment across conditions by comparing the set with differentially expressed genes and transcripts (from DESeq2-based gene-and transcript-level analyses) to distinguish switches accompanied by changes in gene expression from “switch-only” regulation. Selected loci (e.g. PFKFB3, EIF3E) were visualized to illustrate condition-specific isoform usage and predicted structural changes.

